# Central tendency and serial dependence in vestibular path integration

**DOI:** 10.1101/2024.06.25.600645

**Authors:** Sophie C.M.J. Willemsen, Leonie Oostwoud Wijdenes, Robert J. van Beers, Mathieu Koppen, W. Pieter Medendorp

## Abstract

Path integration, the process of updating one’s position using successive self-motion signals, has previously been studied using visual distance reproduction tasks in which optic flow patterns provide information about traveled distance. These studies have reported that reproduced distances show two types of systematic biases: central tendency and serial dependence. In the present study, we investigated whether these biases are also present in vestibular path integration. Participants were seated on a linear motion platform and performed a distance reproduction task in total darkness. The platform first passively moved the participant a pre-defined stimulus distance which they then actively reproduced by steering the platform back the same distance. Stimulus distances were sampled from short- and long-distance probability distributions and presented in either a randomized order or in separate blocks to study the effect of presentation context. Similar to the effects observed in visual path integration, we found that reproduced distances showed an overall positive central tendency effect as well as a positive, attractive serial dependence effect. Furthermore, reproduction behavior was affected by presentation context. These results were mostly consistent with predictions of a Bayesian Kalman-filter model, originally proposed for visual path integration.

**New & Noteworthy:** Distance reproduction tasks based on visual information about the traveled distance have shown that reproductions are biased by central tendency and serial dependence effects. Here, we show that distance reproductions based on vestibular signals show similar biases and that the reproductions are affected by the presentation order of the stimulus distances.

## Introduction

How do we keep track of our position when navigating our surroundings? An important aspect of human spatial navigation is path integration, which is the process of continuously updating one’s position using successive self-motion signals (1, 2). These self-motion signals can come from various senses, such as the visual and vestibular systems (3) and can also be derived from motor signals (4–6).

To investigate the mechanisms underlying path integration, studies often make use of distance reproduction tasks (7–9). Typically in such tasks, a participant is virtually or physically moved an unspecified distance and then asked to reproduce that same distance. Generally, participants show systematic biases in their reproductions. Some studies show an overall overestimation (8, 10) while others report an underestimation of the reproduced distance (11, 12). There is also work that reports that shorter distances are overestimated while longer distances are underestimated (7, 9, 13–18).

The underestimation of traveled distance has been modeled by assuming that the integration of self-motion information is leaky (19, 20). This model can also predict an overestimation if instead of the already traveled distance, the remaining distance to a target position must be judged (20). However, these types of models cannot explain the observation that reproduced distances are also affected by the history of experienced distances (7). Indeed, several studies have shown that path integration is biased by the distribution of distances a participant encounters during an experiment as well as the sequence in which these distances are presented (21, 22). The former is referred to as the central tendency bias (23) and the latter as serial dependence (24, 25) – both well-known observations across perceptual domains (26). What is the origin of these biases in path integration?

Recent studies suggest that the observed biases do not reflect a distorted integration process, but rather arise from probabilistic computations that perform near-optimal Bayesian inference on noisy but unbiased self-motion velocity signals (7, 18, 27–29). In more detail, because sensory information and motor commands, as well as the neural processing itself, are endowed with intrinsic random noise, the self-motion cues should not be treated as point estimates but rather be approached as probability distributions. For path integration, the Bayesian framework states that the observer estimates the most probable distance (the posterior) by integrating noisy sensory signals (the likelihood) with prior expectations (as derived from past experiences), following Bayes’ rule.

In support, Lakshminarasimhan et al. (27) found that the bias in visual path integration was better explained by a Bayesian prior favoring slower speeds than by leaky integration of unbiased self-motion velocity. Glasauer and Shi (22) showed that a static Bayesian prior, i.e., a distribution with a fixed variance and a fixed mean, could account for central tendency biases in visual path integration but could not explain the serial dependence effects showing that responses were attracted towards the previously presented distance. To account for both types of biases, they proposed instead a Bayesian model that assumes that stimuli are drawn from a distribution with a fixed variance but whose mean changes from trial to trial (22). Can this model also explain the biases in vestibular (or more generally, idiothetic) path integration?

In the present study, we first investigated whether the central tendency and serial dependence effects are also observed in vestibular path integration. If path integration relies on a single multimodal representation of estimated distance irrespective of the type of sensory input, then we expect to find similar central tendency and attractive serial dependence biases as observed in visual path integration (22). However, it is also possible that the reproduced distances show no or even repulsive serial dependence, biasing perception away from the previously presented distance to increase overall sensitivity to different distances instead of keeping the continuity of vestibular path integration (21). To obtain more insight into the origin of the biases, we studied if these biases are affected by the presentation context in which the distance stimuli are experienced. To do so, we sampled distances from two probability distributions covering a range of ‘short’ and ‘long’ distances and created two different contexts by changing the order in which the stimuli were presented. Second, we tested whether the reproduced distances and observed biases, and their potential dependence on presentation context, could be explained by the model proposed by Glasauer and Shi (22) for visual path integration.

## Methods

### Participants

Thirty-one participants took part in the experiment. All participants were naïve as to the purpose of the study and had normal or corrected-to-normal vision as well as no hearing issues or history of motion sickness. The experiment took about 90 minutes and participants received course credits or €15 as reimbursement. One participant was excluded due to problems with sound masking during the task, so that data of 30 participants (11 men and 19 women, aged 18-31 yr) are reported. The study was approved by the ethics committee of the Faculty of Social Sciences of Radboud University Nijmegen and all participants gave written informed consent.

### Setup

We implemented a distance reproduction task using a vestibular sled, consisting of a chair mounted on top of a linear motion platform, that moved along the participant’s interaural axis on a ∼95 cm long track (see Figure 1). A linear motor (TB15N; Tecnotion, Almelo, The Netherlands) and servo drive (Kollmorgen S700; Danaher, Washington, DC) were used to power and control the platform. Participants wore a five-point seat belt and their head was fixated using ear cups. The chair contained emergency buttons which could be pressed at any time during the experiment to stop the motion of the platform. The platform could move passively, i.e., outside of the participant’s control, or actively, by the participant rotating a steering wheel (G27 Racing Wheel; Logitech, Lausanne, Switzerland) which was mounted on a table at chest level in front of them. The steering wheel had a range of rotation from -450° to +450° with a resolution of 0.0549° and encoded the linear velocity of the sled (1 cm/s per degree). The task took place in total darkness and did not contain visual stimuli. We used an OLED screen (55EA8809-ZC; LG, Seoul, South Korea) placed in front of the sled to present instruction messages that explained the task before data collection started. The participant wore in-ear headphones with active noise cancelling (QuietComfort 20; Bose, Framingham, Massachusetts) that played a white noise sound to mask noise produced by the motion platform, alternated by single-tone beeps indicating the different stages of each trial. The experiment code was written in Python (v.3.6.9, Python Software Foundation).

**Figure 1.**
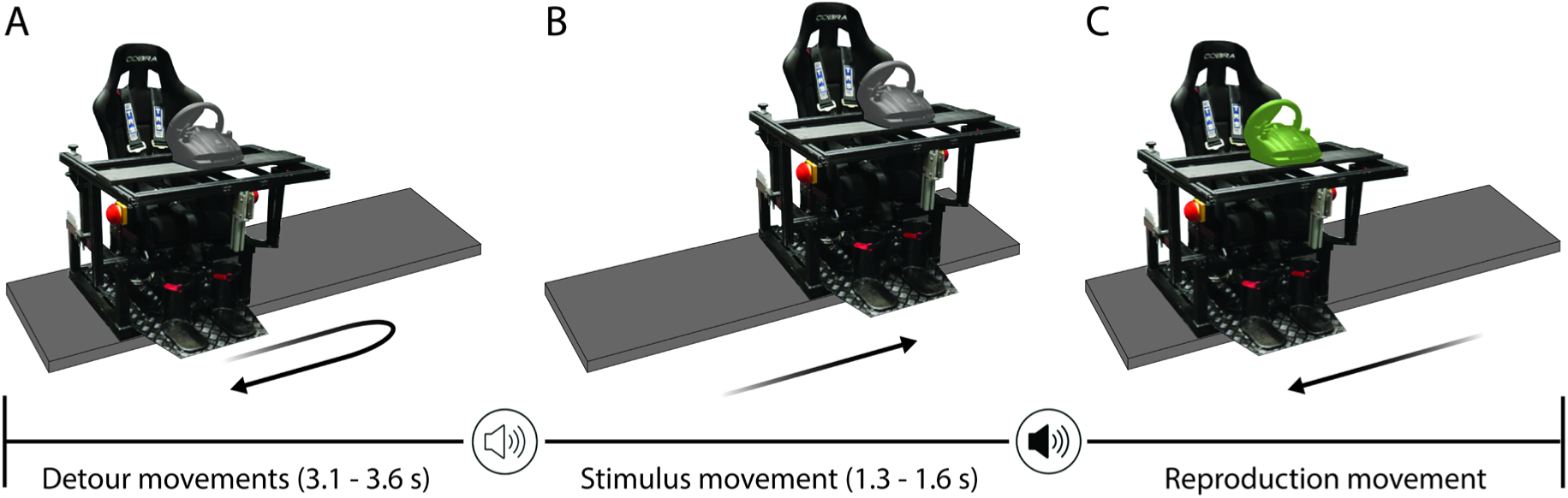
Distance reproduction task in a vestibular sled. Participants were seated in a chair mounted on top of a linear motion platform. At the start of each trial, participants were moved to a new start position via two detour movements (A). Then, a low-tone beep cued that the participant would be moved over the stimulus distance (B). Finally, a high-tone beep instructed the participant to use the steering wheel to move the sled over the same distance in the opposite direction: the reproduced distance (C).

### Reproduction task

Participants performed a vestibular distance reproduction task (see Figure 1). Every trial started with a *stimulus* movement, where the chair was passively moved by a pre-defined distance. The participant’s task was then to actively reproduce the distance by steering the sled in the opposite direction: the *reproduction* movement. In essence, the participant always had to move the chair back to the location from which the stimulus movement started. The direction of the stimulus movement was the same across the trials of one participant. Half of the participants were randomly assigned to leftward stimulus movements and the other half to rightward stimulus movements.

Before the stimulus movement, the chair was passively moved via two random detour movements to one of two start locations, to ensure enough space on the track for the upcoming stimulus movement. Detours were used to prevent participants from receiving feedback about their previously reproduced distance. The first detour moved the chair to a random location within ±30 cm from the middle of the track with a random duration between 1.8 and 2.3 seconds. The second detour movement subsequently brought the chair to the start location in 1.3 seconds. This start location was on the left side of the track for rightward stimulus movements and on the right side of the track for leftward stimulus movements.

Subsequently, a low-tone beep was played to alert the participant to the upcoming stimulus movement. This movement had a random duration between 1.3 and 1.6 seconds. The lower bound was determined such that none of the stimulus movements had a peak acceleration exceeding 1G and a peak absolute velocity exceeding 100 cm/s. The upper bound resulted in the shortest stimulus movement to have a peak acceleration of ∼38 cm/s^2^ and a peak velocity of ∼20 cm/s, such that the vestibular thresholds for perceiving the direction of linear lateral movements were well exceeded (30). All passive movements, i.e. the detours and stimulus movement, followed a minimum jerk profile.

After the stimulus movement finished and a random waiting time between 0.5 and 1 s had passed, a high-tone beep cued the participant to make the reproduction movement by steering the sled in the opposite direction for the same distance as the stimulus movement. If the participant moved the steering wheel too soon (i.e., before the beep), the trial was aborted. The angle of the steering wheel encoded the chair’s linear absolute velocity (1 cm/s per degree) and this gain was kept constant throughout the experiment. Participants could steer the sled up to a maximum absolute velocity of 100 cm/s and could stop the movement by rotating the steering wheel back to the upright position. The chair stopped moving when the absolute velocity became lower than 2 cm/s, after which the trial ended. The movement also ended when the absolute velocity fell below 6 cm/s while the steering wheel angle was constant for 100 ms or the steering changed direction. Participants were instructed to make one smooth movement, i.e., it was not possible to steer back or resume steering after the chair had come to a stop. Participants were free to choose the duration of their reproduction and did not receive feedback about their reproduction performance (except in the training block, see below).

### Paradigm

We sampled the stimulus distances from two probability distributions covering a range of ‘short’ and ‘long’ distances (see Figure 2A). Because magnitudes seem to be internally represented on a logarithmic scale (7, 28, 31–33), we sampled the stimulus distances from log-normal distributions. Log-transforming these distances yielded equal-variance normal distributions (see Figure 2A, inset). Before log-transforming, we divide the distances by a reference distance (1 cm) so all log-transformed distances are dimensionless. Stimulus distances on linear scale varied overall between 17 and 60 cm. The medians of the short and long stimulus distributions were 24.6 cm and 45 cm, respectively, where the distance between the medians of the distributions was determined such that there was a negligible probability of 0.0001 for a random draw from the long distribution to be shorter than a random draw from the short distribution. The variances of the short and long lognormal distributions were 8.2 and 27.4 cm^2^, respectively.

**Figure 2.**
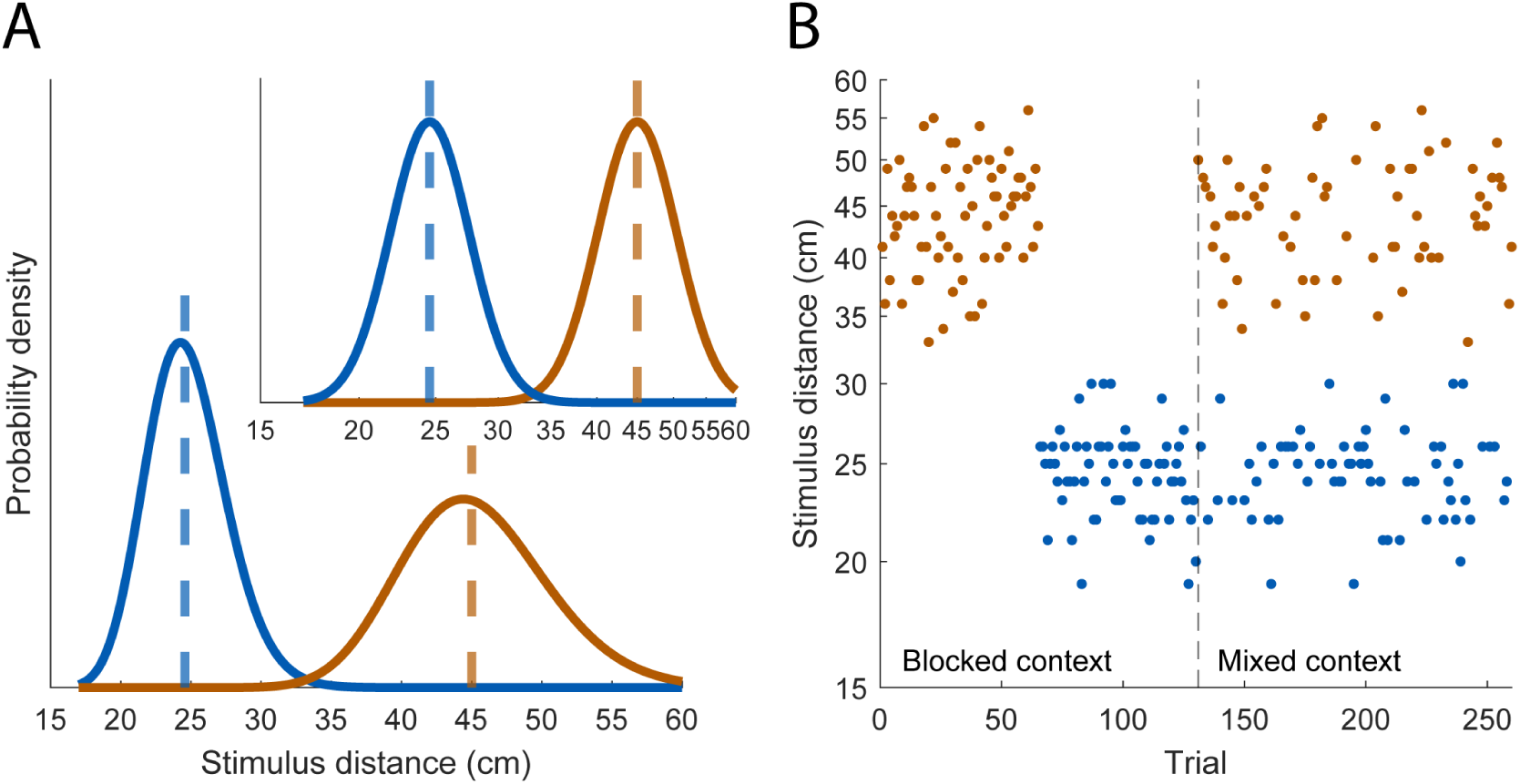
*A*: Distributions of stimulus distances. Distances were sampled from two log-normal probability distributions on linear scale covering a ‘short’ (blue) and ‘long’ range of distances (red). Dashed lines indicate the median distance. *Inset*: The same probability distributions on logarithmic scale. *B*: Example presentation order of stimulus distances. In the blocked context, short and long distances were presented in blocks; in the mixed context, the same distances were randomly interleaved.

Per participant, we randomly sampled 65 distances from each distribution and used these to generate two presentation contexts (see Figure 2B). In the ‘blocked’ context, the short and long distances were presented in separate blocks and therefore separated in time, whereas in the ‘mixed’ context the same short and long distances were randomly interleaved. Participants experienced both contexts during one experimental session of 260 test trials, where the order of the contexts (including the order of the short and long blocks in the blocked context) was counterbalanced across participants. There was no instruction about the existence of the two types of distances and contexts. After every 52 trials (i.e., ∼ 10 min) there was a short break (∼ 2 min) during which the lights were turned on to prevent dark adaptation.

The experimental session started with 20 training trials to familiarize the participant with the task. The stimulus distances in the training trials were drawn from a uniform distribution on linear scale, covering all possible distances (17 to 60 cm). The training trials took place in the dark and followed the same paradigm as the test trials. Contrary to the test trials, the training trials ended with visual feedback on the reproduction error: after the reproduction movement ended, the signed reproduction error in cm was presented on the screen. The training trials were not analyzed.

### Data analysis

Data from the test trials were processed offline in MATLAB (v.R2019a; MathWorks). The end of steering was defined as the first time point with sled speed < 2 cm/s, or as the time point with speed < 8 cm/s when the steering wheel angle remained constant for at least 100 ms or the steering direction changed. The reproduced distance was taken as the distance between the end and start point of the reproduction movement in cm. The reproduction error was defined as the difference between the reproduced and stimulus distance in cm, where negative and positive values represent an undershoot and overshoot, respectively. Trials in which the reproduction movement started too soon, or with reproduction movements in the wrong direction, or with reproduced distances of less than 1 cm were excluded (mean ± SD: 4 ± 4 trials). Because there was no effect of movement direction on the mean unsigned reproduction error across trials (Wilcoxon rank sum test, *p* = 0.300, *r* = 0.19), we regarded all participants as one group.

Central tendency was defined as 1 minus the slope of the linear least-squares regression of the reproduced distance on the stimulus distance on logarithmic scale. We computed the central tendency of the short and long distances separately within each presentation context. We tested whether there was an effect of distance type (short/long) and context (mixed/blocked) on the central tendency values using a repeated-measures ANOVA. Because we found no significant effects (see Results), we averaged the central tendency values for every participant. A one-sample t-test was used to analyze whether central tendencies differed from 0. Partial eta squared 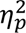 (34) and Cohen’s *d* (35) are reported for the ANOVA and t-test, respectively.

To study if the perception of the short and long stimulus distances was affected by the context in which they were presented, we used the same linear regressions to extrapolate how participants would have reproduced the median distance of the entire distance range (on logarithmic scale, which corresponds to a distance of 31.9 cm on linear scale). We tested the effect of distance type and context on these estimated reproductions using a repeated-measures ANOVA, followed by simple effect tests of the interaction effect levels with Bonferroni correction. Partial eta squared 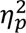 is reported for the ANOVA.

We calculated serial dependence, defined as the slope of the linear least-squares regression of the reproduction error on trial *n* on the stimulus distance on trial *n* – 1 on logarithmic scale (22). The reproduction error was computed by subtracting the reproduced distance on logarithmic scale from the stimulus distance on logarithmic scale. We computed the serial dependence of the short and long distances separately within each presentation context. Because not all of the difference scores were normally distributed, we performed Wilcoxon signed rank tests to analyze whether there were differences in serial dependence values between distance types and contexts. As in the central tendency analysis, we found no significant differences (see Results) and averaged the serial dependence values for every participant. A one-sample t-test was performed to test whether serial dependencies differed from 0. Effect size *r* (36) and Cohen’s *d* are reported for the Wilcoxon and t-test, respectively.

### Modeling

#### The two-state model and special cases

We implemented a Bayesian model, similar to the ‘two-state’ model developed by Glasauer and Shi (22) for visual path integration, to evaluate whether it could also explain the central tendency and serial dependence biases in vestibular path integration. The model first transforms the sensory input *d*_*i*_ to logarithmic scale with *z*_*i*_ = ln(*d*_*i*_), to which the following three generative assumptions are applied:

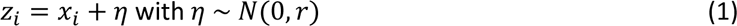

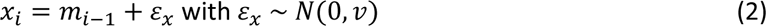

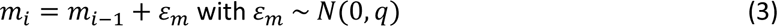

The model thus assumes (i) that the sensory measurement on trial *i*, *z*_*i*_, is drawn from a normal distribution centered on the log-transformed stimulus distance *x*_*i*_ with a fixed variance *r* (Equation 1); (ii) that the stimulus distance *x*_*i*_ is drawn from a normal distribution with mean *m*_*i*−1_ and fixed variance *v* (Equation 2); and (iii) that the mean of this distribution *m*_*i*_ varies over trials following a random walk with a fixed variance *q* (Equation 3). The stimulus distance *x*_*i*_ and the mean of the stimulus distribution *m*_*i*_ are the two states of the two-state model, which are estimated on every trial by a time-discrete Kalman filter. Here, the Kalman filter estimates of the two states on trial 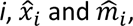 are based on the sensory measurement *z*_*i*_ and the estimated mean of the stimulus distribution on the previous trial 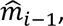 which are weighted by the Kalman gain (see Appendix for the equations). The final estimated reproduced distance on trial *i* on logarithmic scale is computed as 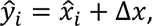 where Δ*x* is a shift term that accounts for global under- or overestimation. In total, the model has four free parameters that are fitted to the reproduction data: the variances *r*, *v*, *q*, and the shift term Δ*x*.

Identical to Glasauer and Shi (22), we considered two special cases of the two-state model based on the assumed stimulus distribution. The ‘static’ variant is obtained by fixing variance *q* at 0, corresponding to a stimulus distribution with a fixed mean. This results in distance estimates that are independent across trials and thus show no serial dependence. In the other special case, the ‘iterative’ variant, variance *v* is set to 0, causing the stimulus distribution to depend on the distance estimate in the previous trial and the estimates to show maximal serial dependence. Both variants, with only 3 free parameters, were fitted to the present data.

#### Sensitivity of the two-state model to different stimulus distributions

The *v* and *q* parameters of the two-state model capture assumptions about the stimulus distribution. To explore to what extent the observed differences in reproduction behavior between presentation contexts can be explained by different assumptions, we adapted the two-state model by introducing separate *v* and *q* parameters for the mixed, short and long blocks. The resulting model has 8 free parameters (*r*, *v_mixed_, v_short_, v_long_, q_mixed_, q_short_, q_long_,* Δ*x*).

We also tested whether the context-dependent differences in the reproduction data are better explained by a block-dependent shift parameter Δ*x*, rather than block-dependent variances. We therefore adapted the two-state model by allowing different shift parameters in the three blocks, while keeping the other parameters constant across blocks, resulting in a model with 6 free parameters (*r*, *v*, *q*, Δ*x*_mixed_, Δ*x*_short_, Δ*x*_long_). Both adapted versions of the two-state model contain only one measurement variance parameter *r* because we assumed that the measurement noise would not change over the course of the experiment.

#### Model fitting and comparison

We determined the log-likelihood of the data given the model parameters across all trials. On every trial, we computed the probability density of the participant’s reproduced distance, given the model’s distribution of possible reproduced distances (equations are included in the Appendix). The free parameters of the models were fitted to the data of each participant individually using the Matlab function *fmincon*, which minimized the total negative log-likelihood summed over trials. Lower bounds were set to 0 for the variance parameters *r*, *v*, and *q*. Start values were set to 1 for the variance parameters and to 0 for the Δ*x* parameter(s) to initialize the GlobalSearch algorithm (Matlab function *GlobalSearch*, (37)), which iteratively executed the *fmincon* function with different start values.

For comparison, we computed the BIC score of each model variant. The BIC is based on the log likelihood, while taking into account the number of free parameters of the model. A lower BIC score indicates a better description of the data. We computed the BIC scores of the model variants relative to the two-state model for every participant separately, which are graphically presented in violin plots showing the median, interquartile range IQR and 1.5x IQR of the relative BIC scores across participants. Median values and IQRs of the fitted parameters across participants are reported because of outliers in the fitted values. To visualize the model predictions, stimulus distances as well as actual and predicted reproduced distances were binned into 10 bins per distance type and per context, separately for each participant. Of these variables, we subsequently computed the means and standard errors across participants per bin.

## Results

We studied vestibular path integration by measuring participants’ performance in a distance reproduction task in the dark and analyzing central tendency and serial dependence biases in the reproduced distances. The stimulus distances were either sampled from a short-distance or long-distance probability distribution and different presentation contexts were created by varying the order in which these distances were presented. In the mixed context, the short and long distances were randomly interleaved, whereas in the blocked context, the same short and long distances were presented in separate blocks.

### Central tendency bias

Figure 3A and C show the raw reproduction data of a representative participant, measured in the mixed and blocked context respectively, plotted as a function of the stimulus distance. The regression lines indicate the central tendency. All slopes are smaller than 1, which corresponds to a positive central tendency. A repeated-measures ANOVA on the central tendency values of all participants indicates that there were no significant differences between distance types (short/long, *p* = 0.105, 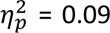) or contexts (mixed/blocked, *p* = 0.245, 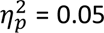), as well as no interaction effect (*p* = 0.091, 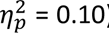). We therefore averaged the central tendency values for every participant and performed a one-sample t-test to study whether the resulting central tendency values differed from 0. On average, we found a positive central tendency effect of 0.39 (SD = 0.21, *p* < 0.0001, Cohen’s *d* = 1.86), which corresponds to a regression line with a slope of 0.61.

**Figure 3.**
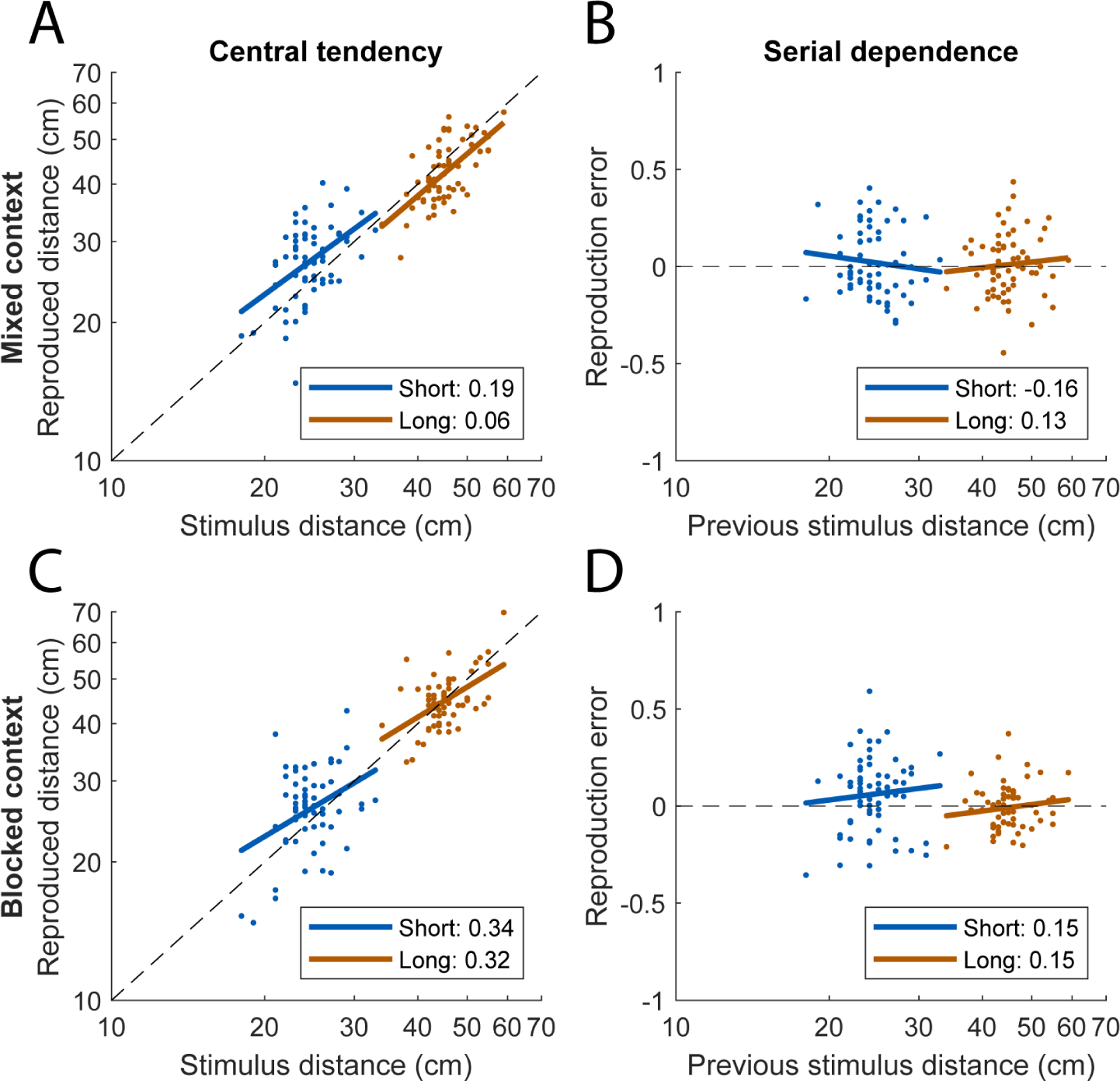
Raw reproduction data of a representative participant in the mixed (A, B) and blocked context (C, D). *A, C*: Reproduced distance as a function of stimulus distance. Regression lines show the central tendency bias. Within each presentation context, separate regressions were performed on the short (blue) and long distances (red). Central tendency values (1 – slope of the regression line) are reported in the legend. *B, D*: Reproduction error on the current trial as a function of stimulus distance on the previous trial, where reproduction errors were computed after transforming the stimulus and reproduced distances to logarithmic scale. Regression lines show the serial dependence bias and were computed for the short and long distances separately. Serial dependence values (slope of the regression line) are reported in the legend.

We performed an additional analysis to study how the short and long stimulus distances were perceived depending on the context in which they were presented. Based on the same linear regressions, we estimated how the median distance of the entire stimulus distance range would have been reproduced in the two presentation contexts (see Figure 4A and B). A repeated-measures ANOVA with factors distance type and presentation context on these estimates, yielded a main effect of distance type, indicating larger estimated reproductions of the median stimulus distance based on the long-distances regression lines than on the short-distances regression lines (*p* = 0.032, 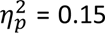). In other words, the median stimulus distance would have been reproduced longer if it had been part of the long as compared to the short stimulus distance range. There was no main effect of presentation context (*p* = 0.563, 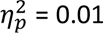), but crucially, there was a significant interaction between distance type and presentation context (*p* = 0.005, 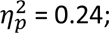 see Figure 4C). Follow-up tests showed that the above-mentioned effect of distance type was only present in the blocked context (mean reproduction based on short-distances regression line = 26.5 cm and long-distances regression line = 29.8 cm, *p* = 0.004), whereas the mixed context showed no difference (mean reproduction based on short-distances regression line = 29.1 cm and long-distances regression line = 28.4 cm*, p* = 0.318). This indicates that reproduction behavior depended on the context in which the stimuli were presented.

**Figure 4.**
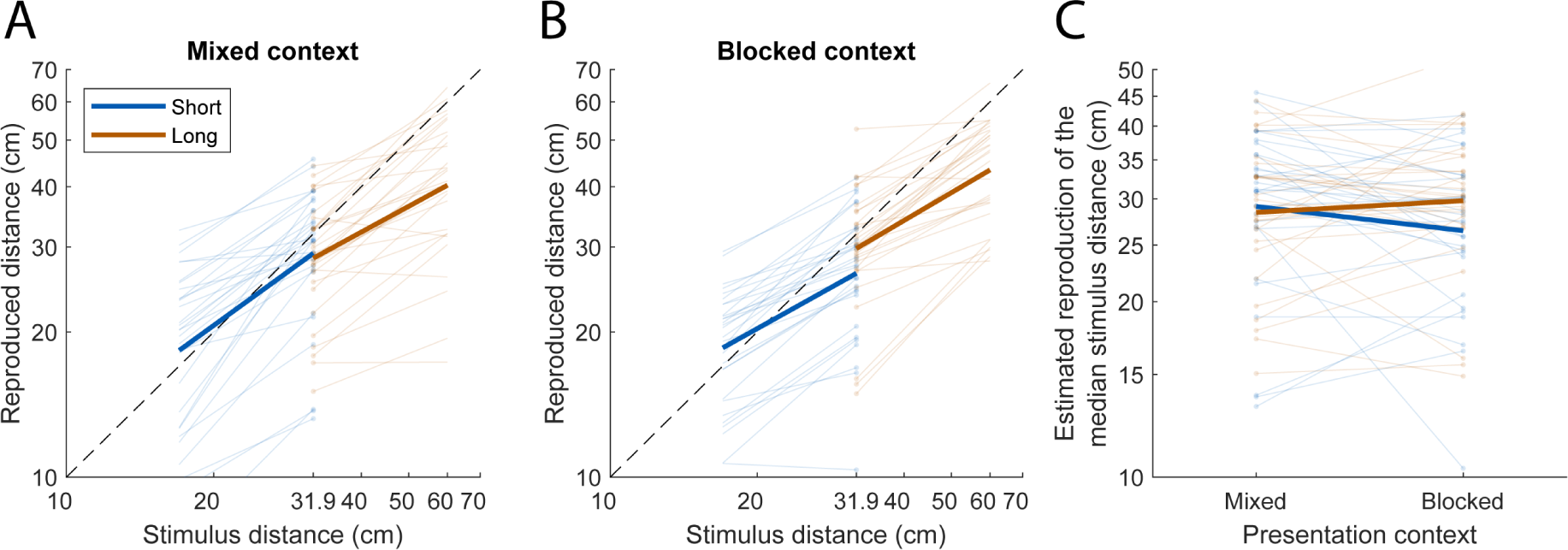
Regression lines based on the stimulus and reproduced distances for short (blue) and long (red) distances in the mixed (A) and blocked context (B) for all participants (transparent lines) as well as the mean (bold lines) on logarithmic scale. The dots represent the estimated reproduced distance at the median of the entire distance range. *C*: Distance-type-by-context interaction effect on the estimated reproductions of the median stimulus distance.

### Serial dependence bias

As an illustration of serial dependence, Figure 3B and D show the reproduction error on trial *n* plotted against the stimulus distance on trial *n* - 1 for the same participant as in panel A and C. The regression lines of the illustrated participant show a slight positive serial dependence, except for the short distances in the mixed context. We performed Wilcoxon signed rank tests on the serial dependence values of all participants and found no significant main or interaction effects for distance type and context (all *p*-values >= 0.185, all *r*-values <= 0.24). After averaging the serial dependence values for every participant, a one-sample t-test revealed a positive serial dependence of 0.13 (SD = 0.15, *p* < 0.0001, Cohen’s *d* = 0.87). This suggests that the reproduced distance on trial *n* is attracted towards the stimulus distance on trial *n* - 1.

### Both the static and two-state models can explain vestibular path integration behavior

We fitted the static, iterative and two-state model variants to all trials of a participant examining a computational, i.e. Bayesian, explanation of these findings. As explained in detail in the Methods, the static model variant assumes that the distance stimuli come from a static, trial-independent stimulus distribution, while the iterative model variant dynamically updates the stimulus distribution on each trial. The two-state model variant combines the static and iterative variants and assumes that the stimulus distance on each trial comes from a distribution with a fixed variance and a mean that can change from trial to trial.

Figure 5 shows the mean binned data and model predictions of the three fitted model variants. In the mixed context, all model predictions are close to the reproduced distance data. In the blocked context, the static and two-state models also provide a relatively good explanation of the reproductions. However, the iterative model underestimates the participants’ reproductions of the short distances and overestimates those of the long distances.

**Figure 5.**
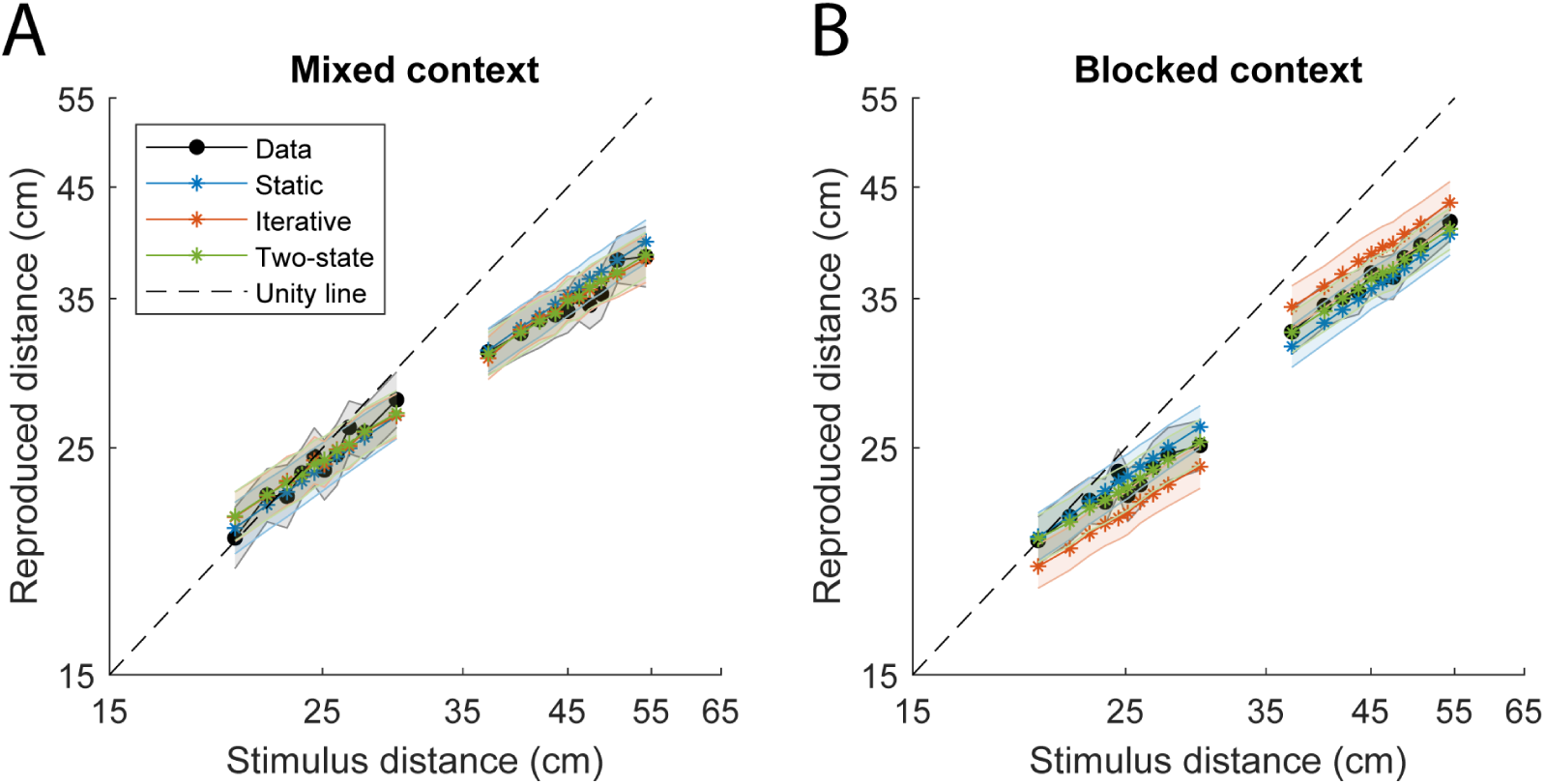
Binned reproduced distances predicted by the static (blue), iterative (red) and two-state model variants (green) as a function of stimulus distance on logarithmic scale for the mixed (A) and blocked context (B). Symbols represent the mean per bin and shaded areas show ± SE, both computed across participants. Unity lines (dashed) show where stimulus and (predicted) reproduced distances are equal.

Next, we computed the central tendency and serial dependence biases as predicted by the three model variants. As illustrated in Figure 6, the predictions of all model variants show relatively similar central tendency values as found in the data. The serial dependence on the other hand is less well predicted by the models. In general, the iterative model overestimates the serial dependence in the reproduction data except for the participants that show a high serial dependence (> 0.30). The static and two-state models tend to underestimate the serial dependence in the data, with the two-state model more often predicting values closer to those found in the actual reproductions than the static model. The figure also shows that the best models in terms of BIC score are also close to the unity line for the central tendency but that this is not the case for the serial dependence.

**Figure 6.**
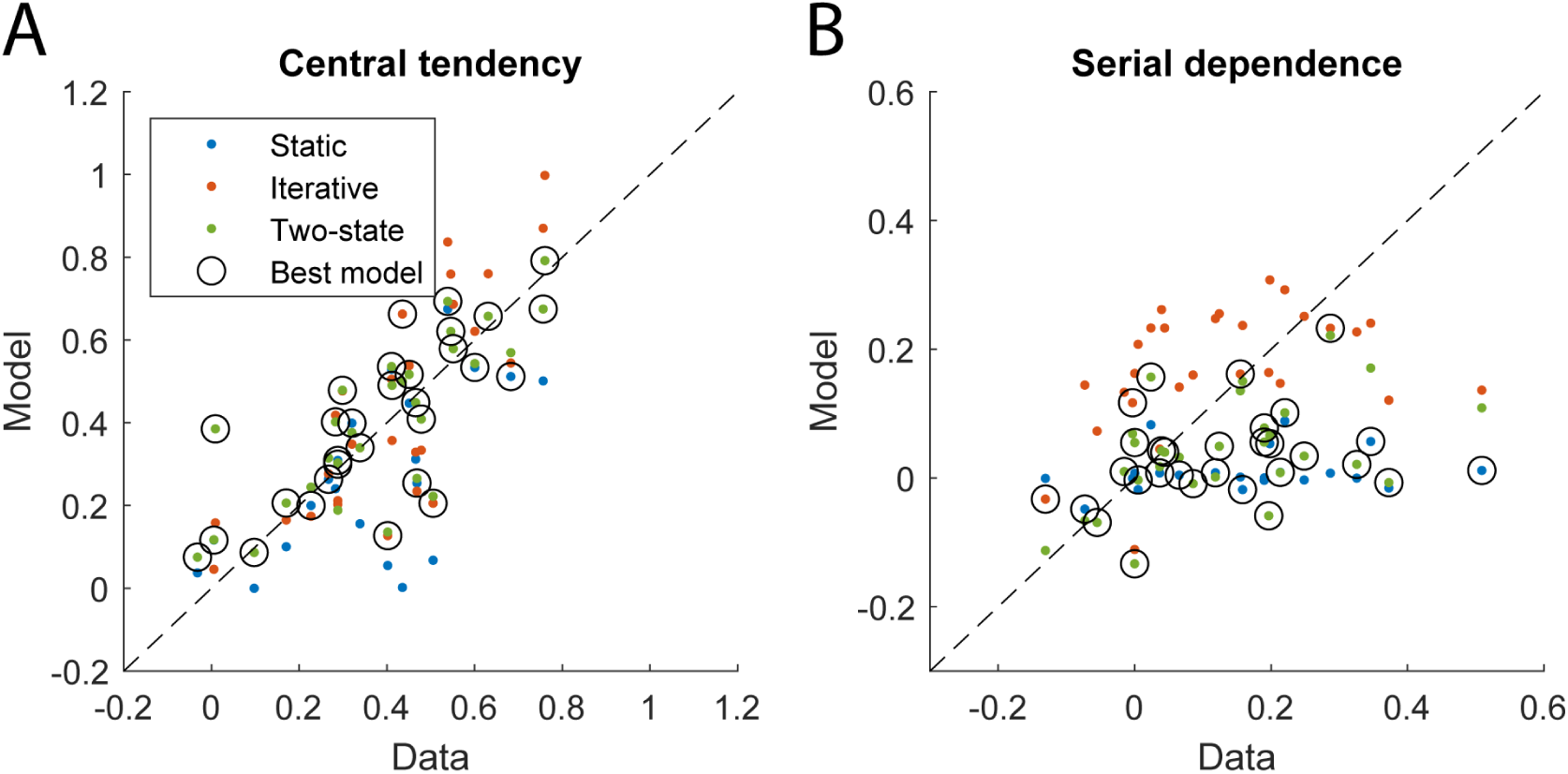
Central tendency (A) and serial dependence values (B) for the static (blue), iterative (red) and two-state model predictions (green) as a function of their measured values. Within a color group, each point represents a participant. Black open circles indicate the models with the lowest BIC score per participant and dashed lines show where the biases in the data and model predictions are equal.

The differences in predicted serial dependence between models are reflected by the fitted model parameter values (see Table 1). The median value of the variance parameter *q*, which determines how much the mean of the assumed stimulus distribution varies over trials, is 0.14 for the iterative model, resulting in serial dependence in the model predictions. The same parameter has a value close to 0 for the two-state model for most participants, predicting virtually no serial dependence and essentially causing the two-state model to behave as the static model. The fitted models show similar measurement variances (*r*) and shift terms (Δ*x*), whereas fitting the static model to the reproduction data resulted in larger values for the stimulus distribution variance (*v*) than in the case of the two-state model. Notably, the fitted parameters show large inter-subject variability.

**Table 1.**
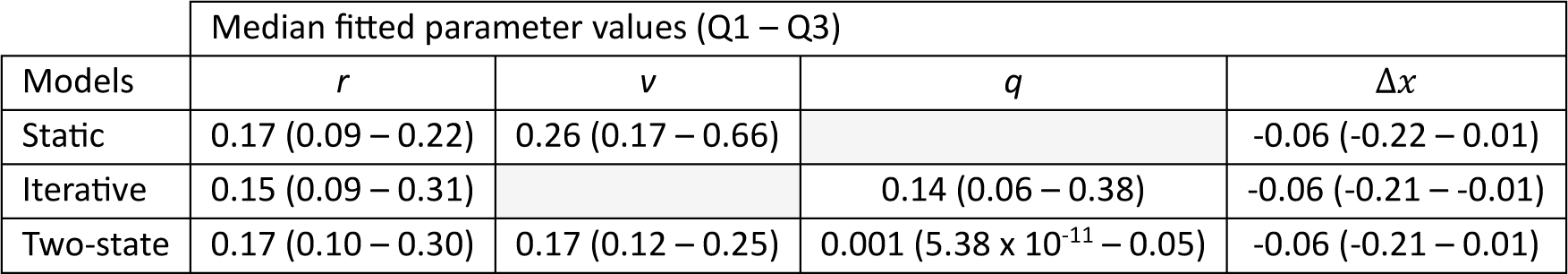
Median (first – third quartile) of the fitted parameter values across participants for the static, iterative and two-state model variants. The parameters *r*, *v* and *q* refer respectively to the measurement variance, the variance of the assumed stimulus distribution and the variance with which the mean of this distribution varies. The Δ*x* parameter is a shift term that represents global under- or overestimation of the reproduced distances.

| Median fitted parameter values (Q1 – Q3) |  |  |  |  |
| --- | --- | --- | --- | --- |
| Models | $r$ | $v$ | $q$ | $\Delta x$ |
| Static | 0.17 (0.09 – 0.22) | 0.26 (0.17 – 0.66) |  | -0.06 (-0.22 – 0.01) |
| Iterative | 0.15 (0.09 – 0.31) |  | 0.14 (0.06 – 0.38) | -0.06 (-0.21 – -0.01) |
| Two-state | 0.17 (0.10 – 0.30) | 0.17 (0.12 – 0.25) | 0.001 ($5.38 \times 10^{-11}$ – 0.05) | -0.06 (-0.21 – 0.01) |

As a final comparison, Figure 7 shows the BIC scores of the static and iterative models relative to the BIC scores of the two-state model, where a relative BIC score > 6 is interpreted as strong evidence in favor of the two-state model (38). Despite substantial interindividual variability, the static and two-state models have similar median BIC scores. The static model has a median relative BIC score of -3.16, describing the data equally well with one less free parameter than the two-state model. The iterative model is outperformed by the two-state model, indicated by a median relative BIC score of 14.16.

**Figure 7.**
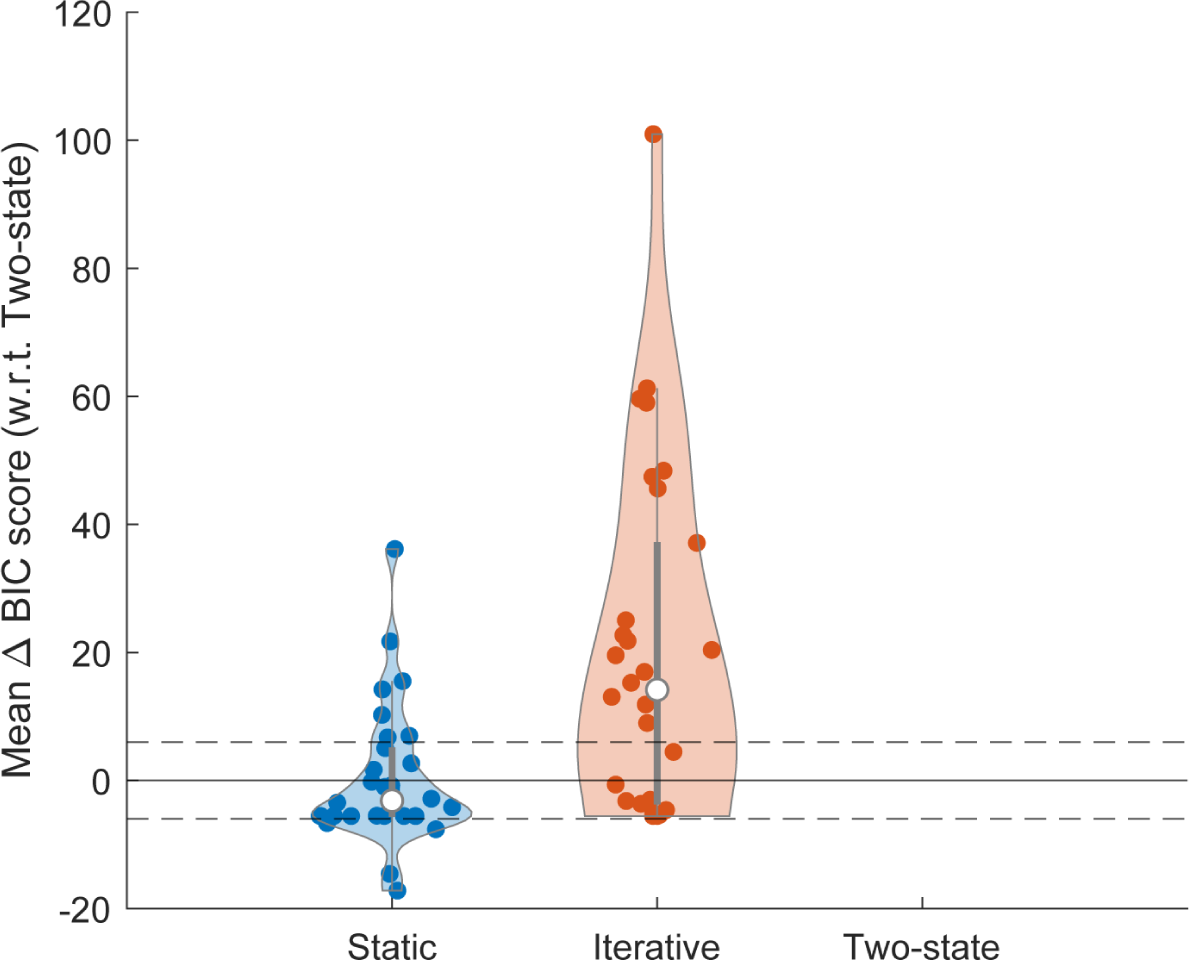
Model BIC scores relative to the BIC scores of the two-state model. Colored data points represent individual participants. White data points show the median relative BIC scores and bold and thin grey lines the IQR and 1.5*IQR across participants. Dashed lines represent a BIC score difference of -6 and 6, where relative BIC scores smaller than -6 or larger than 6 provide strong evidence against or in favor of the two-state model, respectively.

To summarize, the reproduction behavior is best explained by the static and two-state models and less so by the iterative model (see Figure 7), especially in the blocked context (see Figure 5). All models are able to capture the central tendency effects in the data relatively well but perform worse in explaining the serial dependence (see Figure 6).

### The shift parameter of the two-state model can capture context-dependent differences

Can the difference in reproduction behavior across contexts be explained by different assumptions about the experimental stimulus distributions in the different blocks? To examine this, we adapted the two-state model by including separate *v* (representing the variance of the assumed stimulus distribution) and *q* parameters (representing the variance with which the mean of the assumed stimulus distribution changes across trials) for the mixed, short and long blocks (see Methods). The yellow symbols in Figure 8A show the resulting BIC scores relative to the BIC scores of the original two-state model with one *v* and *q* parameter across all blocks. As in the previous BIC score comparison, there is considerable spread in the relative BIC scores. The adapted model had a median relative BIC score of 3.98, thus performing similarly as the original two-state model. In other words, allowing different assumptions about the underlying stimulus distribution between the blocks at the cost of more free parameters does not result in a better description of the vestibular path integration data.

**Figure 8.**
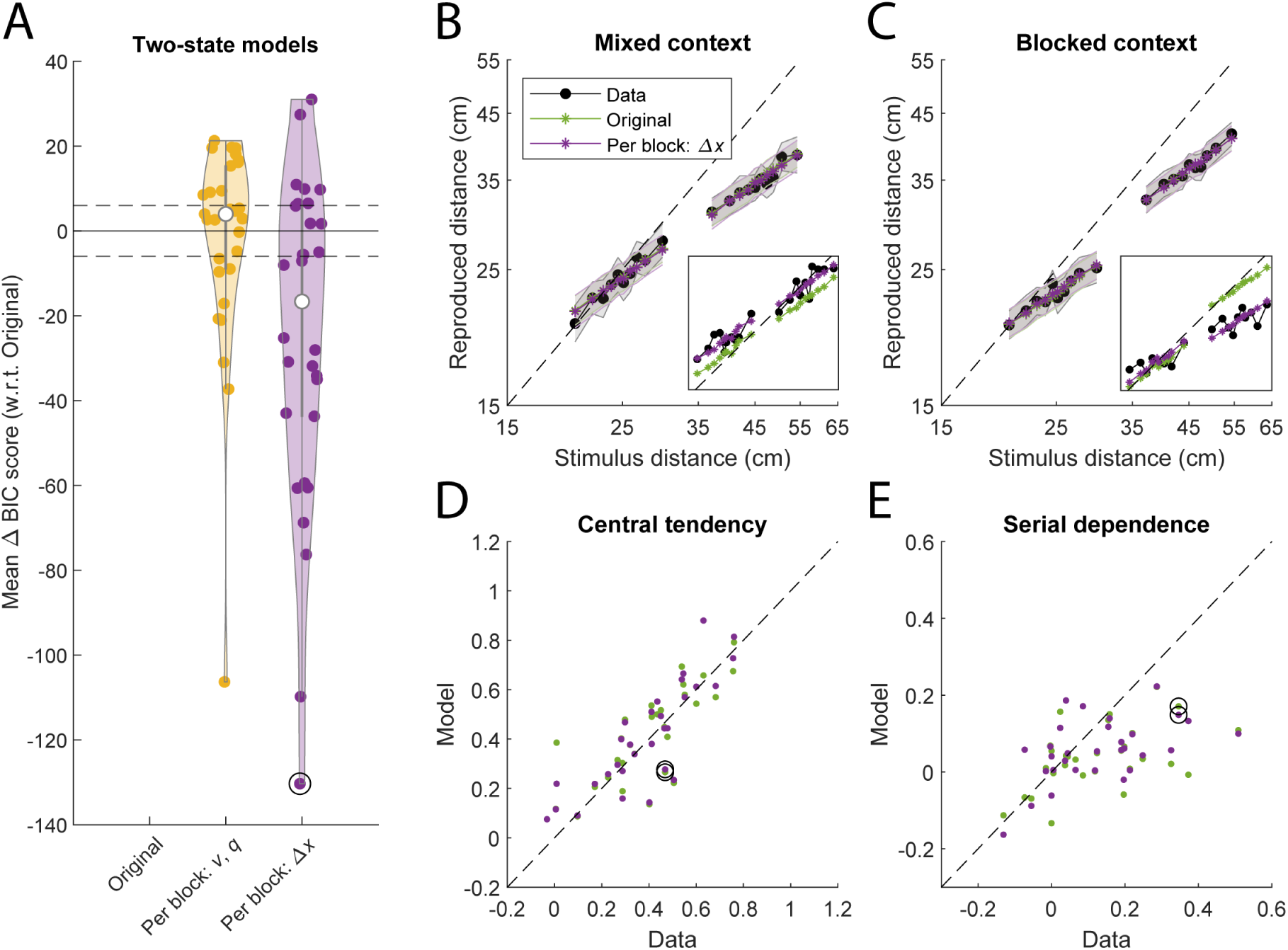
Comparison between original and block-dependent two-state models. *A*: BIC scores of the adapted two-state models with either a free *v* and *q* parameter (yellow) or a free Δ*x* parameter for the mixed, short and long blocks (purple), relative to the BIC scores of the original two-state model. *B, C*: Binned reproduced distances predicted by the original two-state model (green) and the adapted two-state model with a block-specific Δ*x* parameter as a function of stimulus distance for the mixed (B) and blocked context (C). *D, E*: Central tendency (D) and serial dependence values (E) for the same models as a function of their measured values. Panel A, panel B-C and panel D-E are in the same format as in Figure 7, Figure 5 and Figure 6, respectively. Black open circles in panel A, D-E and insets in panel B-C show an individual participant for which the adapted two-state model with block-dependent Δ*x* parameters has the largest decrease in BIC score relative to the original two-state model.

Next, we explored whether allowing separate shift parameters for each of the blocks (Δ*x*_mixed_, Δ*x*_short_, Δ*x*_long_) could explain the differences in the reproductions across contexts. The median relative BIC score of this model with respect to the original two-state model is -16.64 (see Figure 8A in purple), showing that the additional parameters do improve model performance over the original two-state model. This improvement is not directly apparent in the mean model predictions (see Figure 8B and C) or the predicted perceptual biases (see Figure 8D and E), but is visible on the level of the individual participant (see Figure 8B and C, insets). The medians (Q1 – Q3) of the fitted Δ*x*_mixed_, Δ*x*_short_ and Δ*x*_long_ parameter values across participants are -0.08 (-0.21 – 0.05), -0.03 (-0.37 – 0.05) and -0.12 (-0.29 – 0.06), respectively. Taken together, these findings suggest that the differences in reproduction behavior across the contexts are rather explained by block-dependent global underestimations than by block-dependent assumptions about the experimental stimulus distributions.

## Discussion

In this study, we measured human path integration behavior based on vestibular signals and investigated the extent to which distance reproductions show central tendency and serial dependence effects. Participants were seated in a vestibular sled and performed a distance reproduction task in the dark. The sled passively moved the participant with a pre-defined stimulus distance which they actively reproduced by steering the sled back to the location from which the stimulus movement started. Stimulus distances were drawn from a short- and long-distance probability distribution and presented in either a randomized order (the mixed context) or in two separate blocks (the blocked context).

We found a positive central tendency effect that was not affected by distance type (whether the distance was sampled from the short- or long-distance probability distribution) or presentation context (mixed or blocked). The positive central tendency effect indicates that reproductions were drawn towards the mean of the underlying stimulus distribution. This effect has the same direction as the central tendency effects found in visual path integration (7, 14, 22, 29) and other non-visual path integration studies (9, 13, 15–17). This suggests that this bias does not originate within a single sensory modality but might be better understood as the explicit learning of the statistical structure of multimodal motion information.

In addition to a central tendency bias, we found a positive serial dependence bias, again irrespective of distance type or presentation context. This indicates that the reproduced distance on a trial is attracted towards the stimulus distance on the previous trial. This is in line with positive serial dependence effects found in reproductions based on visual information (22). Functionally, positive serial dependence could help to maintain the continuity of the context and promote stable representations for path integration (21, 39).

We implemented a Bayesian model, originally proposed by Glasauer and Shi (22) to explain perceptual biases in visual path integration, to evaluate whether it could also explain vestibular path integration. The model contains three variants based on different assumptions about the stimulus distribution (the static, iterative and two-state variants). On every trial, the model estimates the stimulus distance with a Kalman filter that weights the sensory measurement on the trial with the mean of the stimulus distribution estimated on the previous trial.

We found that the static and two-state model variants provided comparable fits to the vestibular path integration data (see Figure 7). A similar finding was reported by Glasauer and Shi (22), where the two-state model provided the best fit to duration reproduction data for 8 of 14 participants, while for the remaining participants the static model was sufficient. It is important to point out, however, that the model captured the central tendency effect relatively well (see Figure 6) but performed worse in explaining the serial dependence effect. Hence, while the model by Glasauer and Shi (22) provided a joint explanation for the central tendency and serial dependence effects in visual path integration, the present results do not validate this unification in vestibular path integration. Could this be taken to suggest that these biases in vestibular path integration occur due to separate mechanisms?

We prefer to be careful with this conclusion. There are a few differences that should be noted. First, the present data may be noisier, causing the model to not accurately capture all aspects of the reproduction data. Furthermore, the present task was not purely perceptual but also involved a motor component. Compared to a passive reproduction task, additional motor-based self-motion signals could contribute to distance estimates and perhaps trial-to-trial correlations (4, 6, 40, 41). In support, it has been shown that reproducing perceived angular displacements actively reduces the variability compared to passive reproduction (28, 42). The active steering movement in the current task could therefore have introduced non-perceptual effects (i.e., motor biases) in the reproductions. For example, larger distances might have been underestimated more because larger reproduction movements require more effort. For a future experiment, it would be interesting to compare central tendency and serial dependence in active versus passive reproduction tasks of vestibular path integration.

We also studied the effect of different presentation orders on the reproduced distances by presenting the stimulus distances in a mixed and a blocked context (see Figure 2). As indicated by the central tendency analysis, the short- and long-distances regression lines have similar slopes in both presentation contexts. However, the estimated reproductions of the median stimulus distance differ in the blocked context, indicating that presentation context affects vestibular distance reproductions (see Figure 4). A similar interaction between distance type and presentation context was reported in Petzschner et al. (29) for visual path integration, as well as in other magnitude estimation tasks (e.g. Roach et al. (43) for duration reproduction).

To investigate the origin of this interaction, we modified the two-state model by incorporating information about the block (mixed, short, or long) in which the stimulus distance was presented. We first ruled out as a cause of this interaction that participants have different assumptions about the stimulus distribution in the different presentation contexts. More specifically, allowing block-specific parameters for the assumed stimulus distribution (*v* and *q*) did not result in a better description of the reproduced distances (see Figure 8A). Rather, we found that the model variant with block-specific Δ*x* parameters, allowing different global under- or overestimations across blocks, provided a better explanation than the original two-state model. We can only speculate about the functional meaning of this parameter. The global undershooting of reproductions might be caused by increasing uncertainty in the position estimate as more distance is covered (27). Indeed, the observed pattern in the fitted Δ*x*_short_, Δ*x*_mixed_ and Δ*x*_long_ parameter values decreasing with longer distances (-0.03, -0.08 and -0.12, respectively) is consistent with Δ*x* varying linearly with distance, as internally represented on a logarithmic scale. Future studies are needed to examine this potential interpretation.

Furthermore, within this perspective, we emphasize that we studied different presentation contexts, and drew stimuli from a normal distribution rather than a uniform distribution, but we did not vary how we selected the stimuli in each specific block of trials. That is, for each trial, we randomly selected the stimulus distance from the defined stimulus distribution (mixed, short or long). Recently, Glasauer and Shi (44) argued that the central tendency is the result of an unnatural experimental randomization protocol: randomly presenting stimulus distances with large trial-to-trial variability. In many natural circumstances, successive stimuli typically vary only in small range, not randomly jump from one magnitude to another. Using a duration production-reproduction task, Glasauer and Shi (44) showed that the central tendency was greatly reduced if the sequence of the stimulus durations mimicked a random walk compared to that of a randomized sequence. It would therefore be interesting to test how the central tendency effect in vestibular path integration depends on the randomization protocol.

Recent neurophysiological work suggests that the posterior parietal cortex might be involved in producing the perceptual biases affecting path integration. Using a parametric working memory task in rats, Akrami et al. (45) found that the posterior parietal cortex plays a key role in modulating the central tendency bias. When the region was optogenetically inactivated, this bias was not only attenuated, but there was also a suppression of serial dependence, suggesting that the two phenomena may be interrelated. In subsequent neural network modeling work, the same authors explain the two biases through a single mechanism (46). Sensory inputs relayed from the posterior parietal cortex can lead to serial dependence in working memory, from which central tendency naturally emerges.

In conclusion, our results show that distance reproductions based on vestibular signals exhibit positive central tendency and attractive serial dependence, as has been found in visual path integration, suggesting that the biases might arise on a multimodal processing level. Furthermore, reproduced distances were affected by the presentation context of the stimulus distances. The modelling approach suggested that different distance-dependent global underestimations could best account for this contextual effect.

## Appendix

In this Appendix, we provide the equations of the two-state model proposed by Glasauer and Shi (22) and the equations used in the maximum likelihood estimation. The static and iterative model variants are special cases of this model by fixing *q* = 0 or *v* = 0, respectively. (For the definition of parameters *v* and *q*, see the Modeling section in the Methods and below.) Equations 1-3 in the Modeling section can be rewritten in matrix notation as follows:

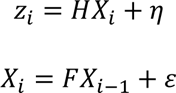

where 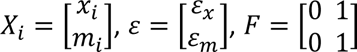 and *H* = [1 0]. The state estimate 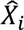 on trial *i* is determined using a time-discrete Kalman filter:

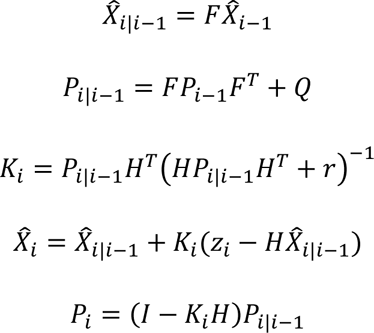

with covariance matrix 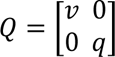 and measurement noise variance *r*. The steady state can be written as:

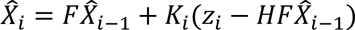

Contrary to Glasauer and Shi (22), we fitted the model’s predictions to the data in logarithmic space using maximum likelihood estimation. Given that there is uncertainty in the measurement *z*_*i*_ on trial *i*, represented by measurement variance *r*, it is possible to compute a distribution of possible reproductions on trial *i*. We computed the expected value 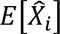 and covariance matrix 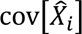 of the estimated stimulus distribution on trial *i* by re-writing the steady-state equation as follows:

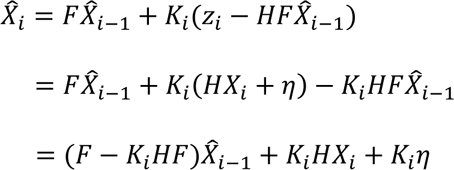

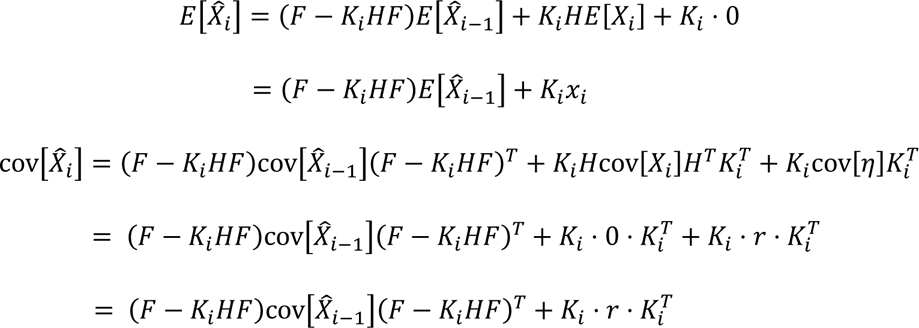

The first element of the resulting expected value vector and covariance matrix correspond to the mean 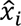 and variance 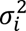 of the estimated stimulus distribution. The model’s prediction of the reproduction on trial *i* is then determined by adding the shift term Δ*x* to the mean of this distribution: 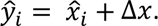 Lastly, the negative log-likelihood on trial *i* (*NLL*_*i*_) is computed based on the probability that the participant’s reproduction on that trial, *y*_*i*_, came from the (normal) estimated response distribution, i.e. the estimated stimulus distribution of which the mean is shifted by Δ*x*:

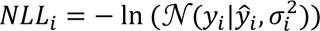

## Acknowledgements

We would like to thank Prof. Dr.-Ing. Stefan Glasauer for the useful discussions about the modeling and their implementation and Dr. James Cooke for his help with the model fitting.

## Author contributions

S.C.M.J.W., L.O.W., R.J.v.B., M.K. and W.P.M. conceived and designed research; S.C.M.J.W. performed experiments; S.C.M.J.W., R.J.v.B. and M.K. analyzed data; S.C.M.J.W., L.O.W., R.J.v.B., M.K. and W.P.M. interpreted results of experiments; S.C.M.J.W. prepared figures; S.C.M.J.W. drafted manuscript; S.C.M.J.W., L.O.W., R.J.v.B., M.K. and W.P.M. edited and revised manuscript; S.C.M.J.W., L.O.W., R.J.v.B., M.K. and W.P.M. approved final version of manuscript.

## Data availability statement

Upon publication, all data and code will be made publicly available on the Radboud Data Repository via the DOI reserved for this collection: https://doi.org/10.34973/rgry-c088.

